# The Movie After-Effect: widespread adaptation of human cortex following naturalistic sensory experience

**DOI:** 10.64898/2026.08.13.744606

**Authors:** Erez Simony, Niv Yahav, Rafael Malach

## Abstract

Neural systems adapt to prolonged sensory input through mechanisms such as gain control or homeostatic plasticity to maintain stable operating ranges. However, this phenomenon has so far been documented under extreme, non-ecological stimuli targeting specific sensory systems. Here, we reveal a widespread adaptation process across diverse human cortical regions following naturalistic movie watching conditions. The effect was evident in 218 out of 251 cortical regions (87%) that exhibit significant stimulus-driven activations. Analyzing the HCP fMRI data set in which 170 participants watched 14 movie clips, followed by rest periods – we found robust evidence for a movie-induced adaptation process, revealed in the post-movies rest periods. Within a region, voxels highly activated at the end of movies subsequently reduced their activity below baseline during rest, with the magnitude of this drop proportional to initial activation levels, manifested as a consistent voxel-population *inversion effect*. Conversely, persistently movie-inactivated voxels exhibited increased activation above baseline. Importantly, this Movie After-Effect (MvAE) enabled successful decoding of the specific rest periods following individual movie clips. Our findings suggest that under naturalistic conditions, cortical neurons dynamically change their gain to achieve homeostatic balance in a process akin to batch instance normalization in artificial neural networks. Whether this ubiquitous MvAE has additional cognitive and memory-related implications remains to be explored.

## Introduction

> *“The living organism is stable. It must be so in order to resist the disruptive forces of the outer world. But it is stable not because it is unchanging, but because it is capable of adjusting itself to change.” — Ludwig von Bertalanffy, General System Theory*

Neural systems must continuously adjust their sensitivity to operate within a stable dynamic range despite large fluctuations in sensory input. A wide range of studies have shown that neurons adapt to sustained stimulation through mechanisms such as gain control, activity rescaling and homeostatic plasticity. These mechanisms are thought to prevent saturation of neural responses and maintain efficient coding of sensory information.

However, most evidence for such adaptation processes comes from highly controlled laboratory experiments that use artificial and often extreme stimuli. It therefore remains unclear whether similar regulatory mechanisms operate during a more naturalistic sensory experience, where stimulation is complex, dynamic and continuously evolving over time.

Indeed, examining the extensive body of research on neuronal adaptation reveals that it has largely focused on artificial laboratory conditions, for example inducing persistently high localized activations^1–3^. Complementing this effect, inducing persistent silencing of cortical neurons leads to increased sensitivity^3^. This bidirectional plasticity manifests in perceptual illusions such as the motion and color aftereffects ^4–6^ and tilt aftereffect^7^. Finally, animal studies combining electrical recordings with fMRI or utilizing intracellular recordings have demonstrated that prolonged activation leads to a decrease in neuronal activity below spontaneous baseline and cellular hyperpolarization^8–10^. The functional role of such effects is most likely homeostasis, aimed at recalibrating neuronal sensitivities to avoid saturation or lower-bound saturation^11,12^. A related function involves internal readjustment of system parameters in response to changing external conditions, such as in prismatic adaptation^13,14^ or norm-based encoding^15^. Typically, the time course of the adaptation effect is rather long requiring many seconds for its induction. In this respect, it should be clearly distinguished from the much faster rapid adaptation effects^16^, which develop over fractions of seconds. Similarly, it should be distinguished from repetition suppression effects, which occur within seconds^17,18^, and likely play a role in e.g. priming effects^19^

While neuronal mechanisms of adaptation have been studied in animal models, much less is known about its neuronal manifestations in humans. Using functional MRI, Tootell et al demonstrated a neural correlation of the perceptual motion after effect in area MT^6^, corresponding to the well-known emergence of an opposite direction of motion sensation following prolonged exposure of a moving stimulus, termed the motion after-effect. However, this result was confined to highly artificial and specific stimulus conditions - and see^20^. While these studies have convincingly demonstrated adaptation effects under well controlled, artificial, laboratory conditions- they leave open the crucial issue- do such effects broadly modulate brain sensitivity during naturalistic, ecological conditions?

In recent years there has been a growing success in extending human neuroscience research into more ecological conditions ^21–23^. In this context, it is of interest to explore whether adaptation effects, mainly studied using artificial stimuli, will be revealed also under naturalistic conditions. To that end we have examined the extensive fMRI data obtained from the human connectome project ^24,25^ – specifically data of human subjects that watched different movie clips.

Our results reveal that during naturalistic conditions, the human cortex shows a significant and widespread adaptation process. Specifically- the effect was robustly revealed through MVPA classification ^26^ during the rest periods many seconds after the movie clip-activations ended. Remarkably, the effect was observed across a wide range of cortical regions including low and high visual areas as well as attention and some of the DMN regions. Thus, our study reveals that during naturalistic stimulation, the gain of human cortical neurons is not stable but undergoes a widespread, dynamic, activity-dependent adaptation. The possible cognitive and functional implications of this wide- spread effect are yet to be discovered.

## Results

Our study was based on the HCP fMRI data^24,25^ which included fMRI scans conducted at 7T in 184 subjects. In the present study we focused on fMRI imaging of brain responses to the presentation of 14 unique audio-visual movie clips from 170 subjects. The structure of the experiment and the data analysis procedures are illustrated in figure 1. Briefly, Functional runs of interest included movie-watching runs followed by resting-state runs. We focused our analysis on two sessions that contained 4 movie-runs each. Each movie-run included 4-5 separate movie-clips. We discarded the last repeated clip from each movie-run and analyzed 14 movie-clips driven BOLD responses and their 14 post-clips resting state periods of 20 sec each. Our analysis included MVPA Classification during movie-clips (Figure 1A), during post-clip resting-states intervals (Figure 1B), and correlation of voxels activity patterns between the end of clips and the end of resting-state periods (Figure 1C). In addition, as a control, we analyzed pure resting-state data (with eyes open with relaxed fixation on a projected bright crosshair on a dark background) from 4 sessions, subdivided into the exact time-periods of the movie-clips and their following resting-state periods, and repeated the same analysis. For further details on imaging data acquisition and analysis, see Methods.

**Figure 1:**
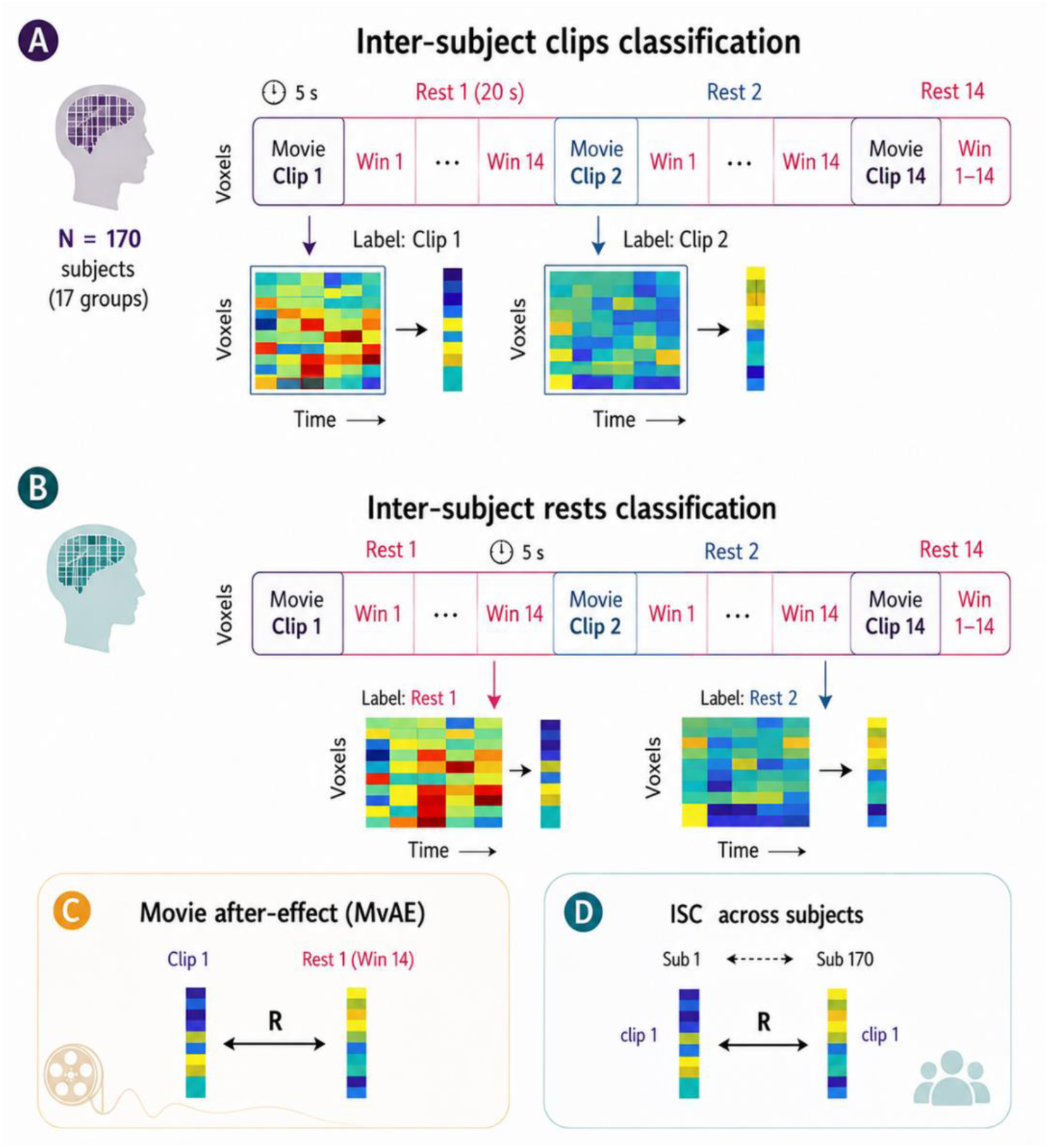
Classification across subjects and the Movie After-Effect (MvAE). During 7T fMRI, participants (N = 170) viewed 14 movie clips interleaved with 20-second resting-state intervals across four functional runs. **(A)** The final 5-second interval of each clip was extracted (voxels x time) from each brain region and averaged over time to yield a spatial voxel pattern per clip. Clip identity was decoded across 300 brain regions using a linear support vector machine (SVM) classifier via leave-one-group-out cross-validation across 17 independent groups (each ‘super-subject’ represents an average of 10 participants). **(B)** Post-clip resting-state intervals were segmented into 14 overlapping 5-second windows. For each window (win1–win14), spatiotemporal activity within a given brain region was extracted and averaged over time. Voxel patterns from the most remote window (win14) were used to train and test an SVM classifier across the 17 super-subjects using a leave-one-group-out framework across all 300 regions. **(C)** The MvAE value was calculated by computing the Pearson correlation between the voxel pattern at the end of a movie clip and the voxel pattern from the final resting-state window (win14). The magnitude of this correlation provides an estimate of the MvAE per clip. **(D)** The spatial Inter-Subject Correlation (ISC) was calculated per movie clip by correlating the end-of-clip voxel patterns across all pairs of the 17 super-subjects.

### Tracing long-lasting adaptation effects

To trace the long-lasting effect of adaptation- we took advantage of the fact that after each movie clip subjects underwent a rest period lasting 20 seconds. In those cases where different movie clips induced different activation levels across cortical regions and voxels- we hypothesized that such differential activations would result in different adaptation across the voxels- and these differences will persist during the resting-state period following each movie clip.

To examine this possibility, we first sought to determine whether the clips elicited sufficiently distinct voxel activation patterns to enable accurate decoding of clip identity based on multivariate activity patterns. Figure 2A depicts the results of these 14-clips support vector Machine (SVM, see methods for details) classification accuracy across 17 groups of subjects, for the entire set of 300 cortical regions. This was done by grouping the data to 17 groups of 10 subjects each. Each group was averaged, yielding 17 “super-subjects”. for each super-subject the last 5-sec from each movie clip was extracted and averaged over time to get a voxels’ activation pattern. These were input to SVM classification, using leave-one-out-cross group validation across 17 super-subjects (for illustration - see Fig 1A). As can be seen, the voxel activations allowed a highly significant decoding of clip identity- ranging from 10% to 95% accuracy across the entire cortex (accuracy threshold = 15%, resting-state-derived null distribution, p < 0.01). In the figure, the cortical regions are arranged in a descending order of their successful decoding, while the colors depict the different networks ^27^.

**Figure 2:**
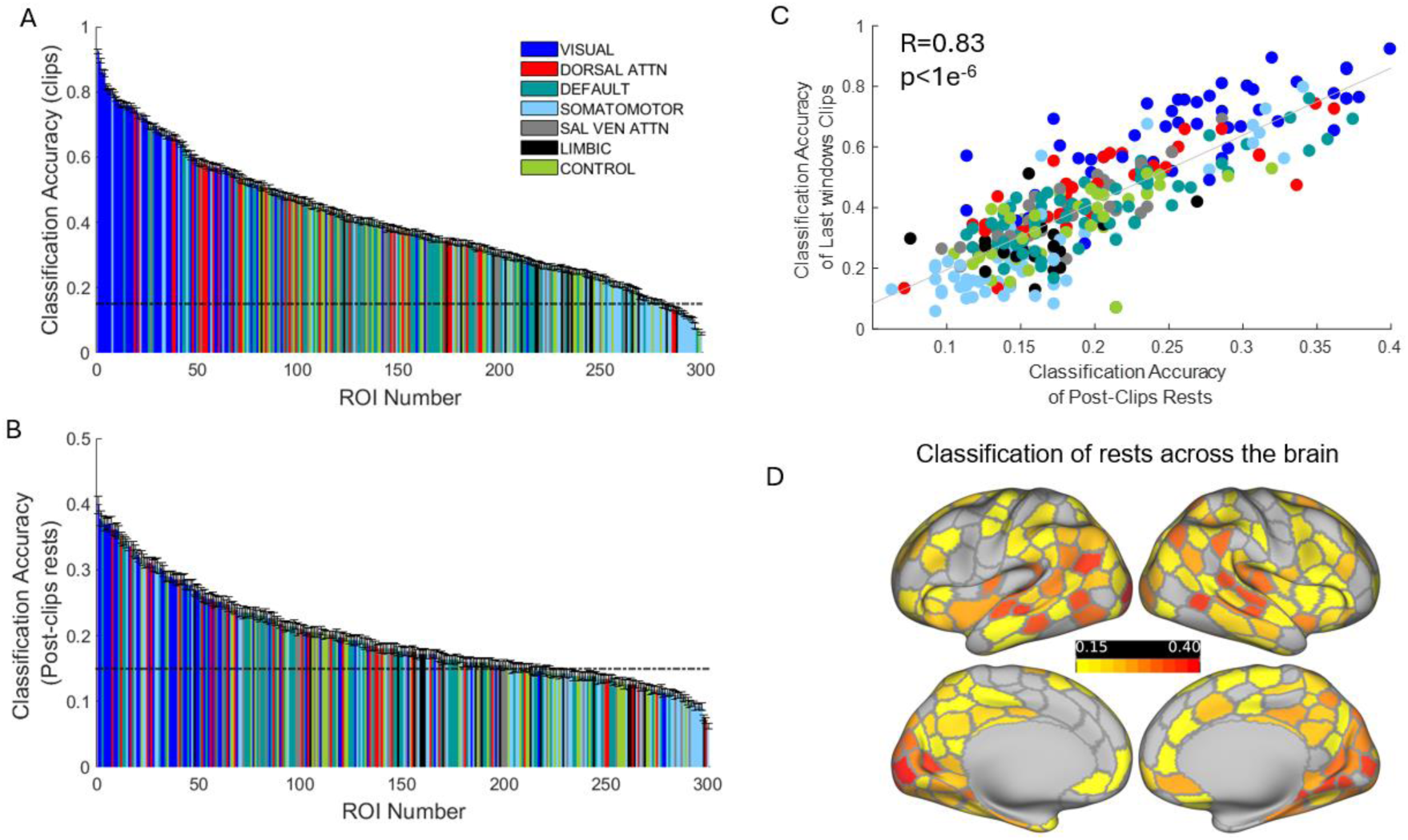
MVPA classification across subjects and 300 brain regions. **(A)** Classification accuracies of the 14 movie clips (mean ± SD across 17 independent groups) using the final 5 seconds of clip data, ordered from highest to lowest decoding accuracy (dashed line indicates the empirical significance threshold at 15%). **(B)** Classification accuracies of the 14 rest periods (mean ± SD across 17 independent groups) evaluated using the final 5-second window of the post-clip resting-state intervals. **(C)** Scatter plot comparing clip classification accuracies against post-clip rest classification accuracies across all regions, demonstrating a highly significant correlation (R = 0.83, p < 10^-6^). **(D)** Surface projection of the classification accuracies during the post-clip resting-state intervals (from panel B) mapped onto the cortical mantle using the 300-region Schaefer parcellation^27^.

Critically, if differential voxel responses to the movies produced a long-lasting trace, it should be possible to decode the identity of the subsequent resting-state periods. To test this, we applied a similar decoding procedure to that used at the end of the clips, but here focused on the rest periods following each movie clip (Fig 1B). To avoid any possible interference from the slow BOLD signal of movie activations - we chose, for classification, the last 5 seconds of the rest period- which was a window separated by 15-20 seconds from the end of the movie clips.

The results of this decoding procedure are depicted in figure 2B with the cortical regions arranged in descending order as in figure 2A. As can be seen, there was a highly significant decoding, solely based on the rest periods across most cortical regions (accuracy threshold = 15%, resting-state-derived null distribution, p < 0.01). Importantly, brain regions that exhibited higher classification accuracy – of the specific clips-towards the end of clips tended to show similarly higher classification accuracy – of the specific rest periods- during the end of the post-clips rests intervals (R=0.83, p<10^-6^, Figure 2C). As depicted in the color coding of figure 2D, the classification accuracies of the last windows of resting-state intervals were widespread across the entire cortex, mainly distributed across the visual, attention, auditory-language networks and some of the DMN regions.

To examine the dynamics of changes in neural activation underlying our ability to decode the resting periods following each movie clip, we tracked the BOLD signal changes of individual voxels (averaged across 170 participants) from the end of the movie clips through the end of the subsequent rest period. Figure 3 shows three examples of such dynamics from three different cortical regions belonging to different networks, and one example from the pure resting-state control. The plots present the neural activation dynamics of individual voxels following the end of a single movie clip. In these plots, voxels are color-coded (from warm to cool colors) according to their activation levels (percent signal change) during the final 5-second period of the movie clip. This analysis revealed a striking population “inversion” effect (Figure 3 A,B) whereby highly activated voxels at the end of the movie clip, from the visual, dorsal attention networks, respectively, reduced their activity below the average baseline activations proportional to the magnitude of activity at the end of clips (the baseline activity is the average activity of 20-sec rest run at the beginning of a movie run). By contrast, voxels from Posterior Cingulate Cortex (PCC; DMN region) were inactivated at the end of the movie-clip and increased their activity above baseline during the post-clip resting interval. The increase in magnitude was proportional to the deactivation magnitude of the voxel’s activity at the end of clips (Figure 3C). Hence, the correlation value between the voxel pattern from the end of clip with the voxel pattern from the end of rest exhibit strong negative correlations (R=-0.82, p<10^-4^, R=-0.79, p<10^-4^, R=-0.47, p<10^-3^). Importantly, resting-state control activity that was virtually segmented similarly to the movie scan, is depicted in Figure 3D. It shows an absence of the inversion effect- so that the average activity of a voxel remained stable over time, resulting in positive correlations between voxel patterns (R=0.2, p<0.05).

**Figure 3:**
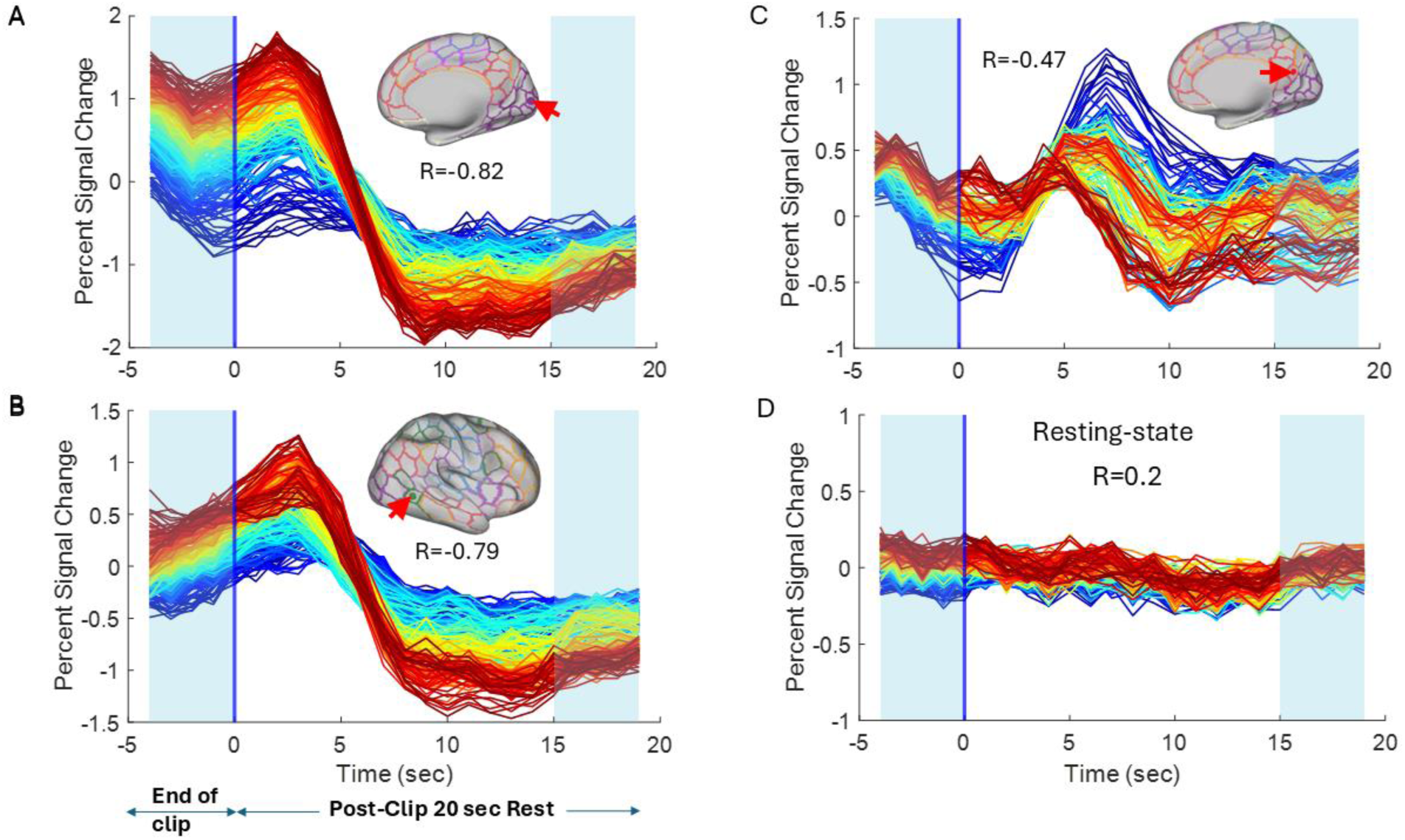
The Movie After-Effect (MvAE) induced population inversion effect. **(A)** Average voxel activity across 170 participants within an early visual area (indicated by the red arrow) during the final 5 seconds of a movie clip (shaded area) followed by a 20-second resting-state interval. Individual voxels are color-coded (from hot to cold colors) according to their activation levels (percent signal change) during the final 5 seconds of the movie clip. The magnitude of the post-stimulus signal drop during rest is proportional to the initial activation level at the end of the clip, resulting in a significant negative correlation (R = −0.82, p < 10^-4^) between the voxel activation patterns at the end of the clip (shaded area) and those at the end of the rest interval (post-clip shaded area). **(B)** A similar population inversion effect is shown for a region within the dorsal attention network (red arrow), demonstrating a significant negative correlation (R = −0.79, p < 10^-4^). **(C)** A region within the Default Mode Network (PCC) displays inverse neural dynamics compared to panels A and B, yet exhibits the same homeostatic inversion effect: the increase in activation during the rest interval is proportional to the deactivation level at the end of the clip, yielding a significant negative correlation (R = −0.47, p<10^-3^). **(D)** Pure resting-state control activity from a representative brain region shows a weak positive correlation between the corresponding virtual clip-end and rest-end periods (R = 0.2, p < 0.05), indicating that spontaneous activity across the 170 participants maintains a baseline positive correlation in the absence of stimulation. Note the consistent inversion of the voxel color hierarchy from the end of the clip to the end of the rest interval induced specifically by naturalistic stimulation.

How consistent was the effect across different movie clips? Figure 4 shows an example of the relation between end of movie-clip voxel activations and end of rest voxel activations in a single area across all 14 movie clips. panel 4A demonstrates all-clips scatter diagram plotting the individual voxel activations at the end of the clip (x-axis) against the activation at the end of rest (y-axis) across all movie clips – depicted in different colors. The overall negative trend of the correlation is evident (R=-0.8, p<10^-4^). Examining the correlations separately across 14 different movie-clips revealed a substantial variability in the anti-correlation strength. This is illustrated in panel 4B, which presents scatter plots for individual movie clips arranged from the most strongly anti-correlated to the least. As can be seen, although the majority of clips produced a robust anti-correlated effect, some clips failed to elicit this pattern. Closer inspection of these scatter plots reveals that a common characteristic of these clips was weak activation at the end of the movie clip, reflected by the clustering of voxels toward the left side of the plots.

**Figure 4:**
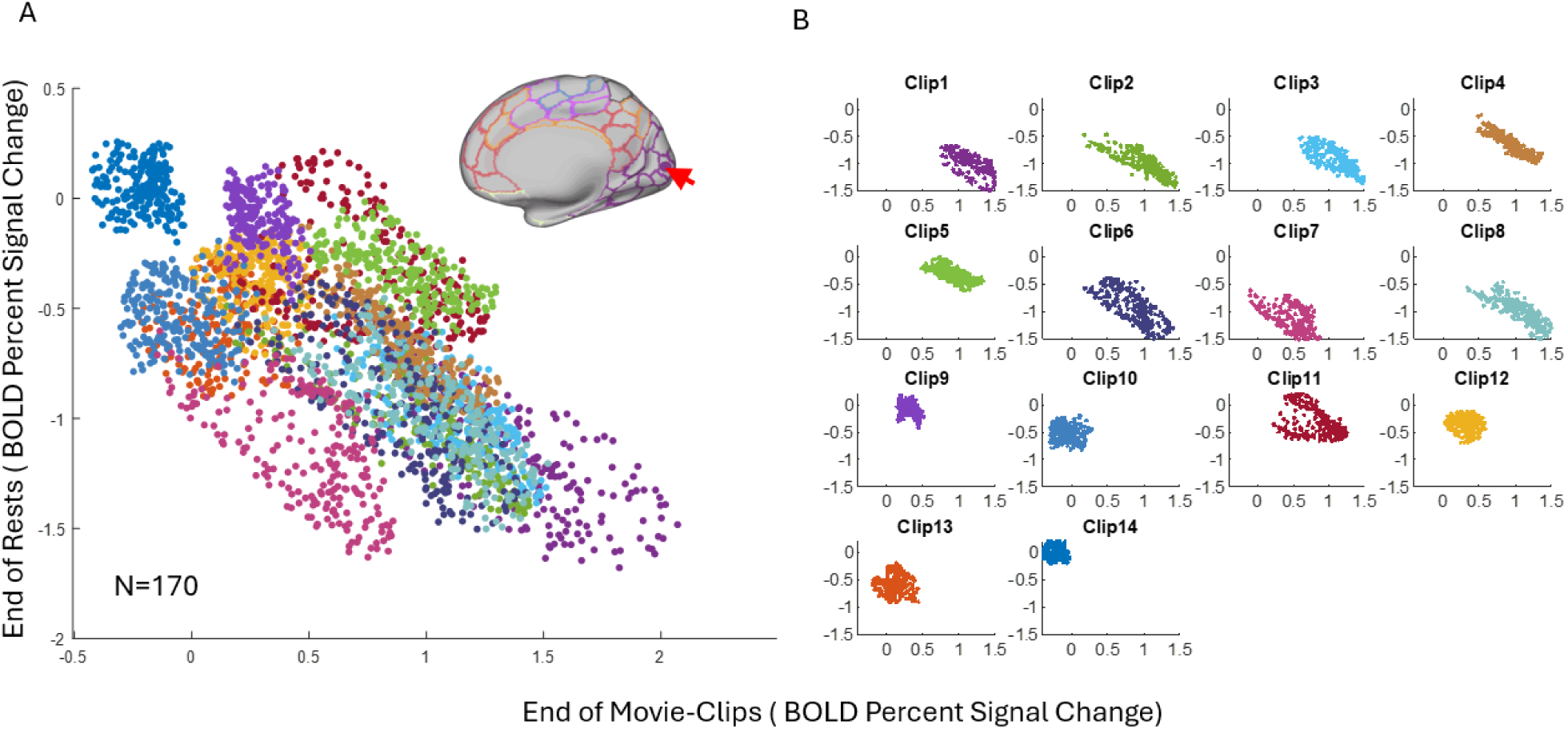
Clip-dependent MvAE. **(A)** Scatter plot displaying voxel-wise BOLD activity (percent signal change) at the end of the rest intervals as a function of activity at the end of the movie clips within a representative visual region (indicated by the red arrow) across all 14 movie clips. Each data point represents the activity of a single voxel, with individual clips depicted in distinct colors. **(B)** Voxel-wise scatter plots separated by individual movie clip, ordered sequentially from the strongest negative inversion effect (highly anti-correlated) to no inversion effect. Note that the magnitude of the negative correlation is tightly coupled to the overall level of regional activation elicited at the end of the clips.

Finally, we examined two aspects of the population inversion effect. First, to what extent differences in the anticorrelation effect across different ROIs, could be explained by the activation of these regions to the movie clips? Second, how widespread was the effect- i.e. to what extent was this anticorrelation significant and widespread across many cortical areas? Figure 5 shows the results of an analysis aimed at addressing these questions. In Panel A, we employed inter-subject correlation (ISC)¹⁸ as an indirect measure of stimulus-induced movie activations. ISC was calculated for each brain region and each clip using the last 5 seconds of each movie clip, and was then plotted as a function of the MvAE for the same clip and region. The strength of the MvAE was defined as follows: the correlation value was calculated for each brain region, per group of subjects (17 groups of 10 subjects each), per clip, by obtaining voxel patterns from the end of one movie-clip (1 x number of voxels per region) and correlating it with the voxels-patterns from the last window of the rest intervals following that clip (Figure 1A, win14). Hence, MvAE value is an overall measure for the adaptation effect, calculated across 14 movie-clips. We found a significant negative correlation between the ISC *per movie-clip* for each brain region and the MvAE value of the same clip, and the same region (R=-0.61, p<10^-5^, see methods for MvAE and ISC calculations).

**Figure 5:**
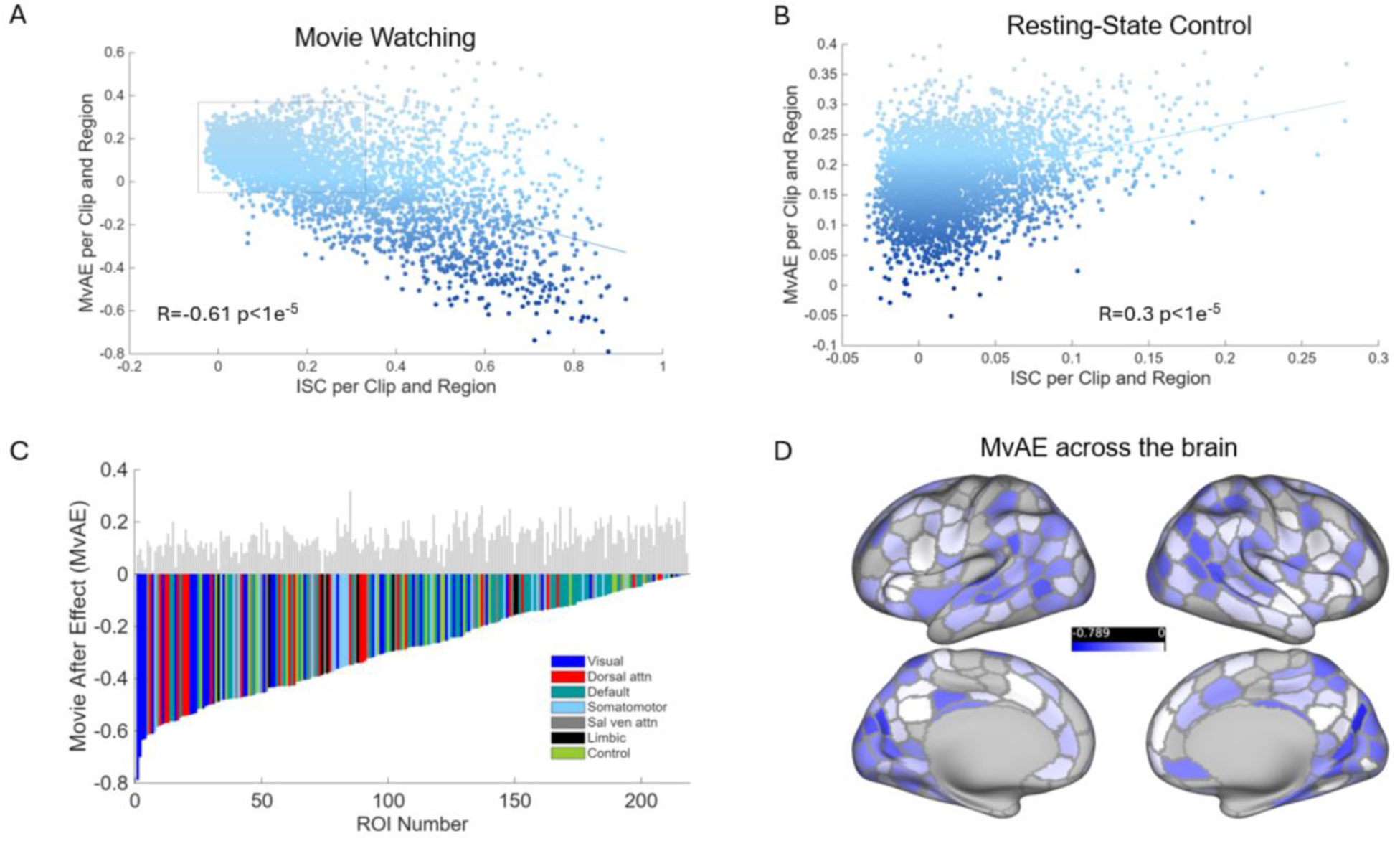
MvAE and ISC values per region and clip. **(A)** Inter-subject spatial pattern correlation (ISpC) calculated at the end of each movie clip across all 300 brain regions (14 clips x 300 regions) is negatively correlated with the corresponding MvAE values (computed as the Pearson correlation between the voxel pattern at the end of a clip and the pattern at the end of the subsequent rest period). **(B)** Identical analysis as in panel A but conducted on pure resting-state control data from the same participants, demonstrating a weak positive correlation between ISC and MvAE per clip and region (note the difference in axis scales, corresponding to the inset dashed rectangle in panel A). **(C)** Widespread distribution of significant negative MvAE values (evident for at least one movie clip per region) across 218 out of 251 brain regions (87%) that exhibited significant inter-subject pattern correlation (R* > 0.25, FWER-corrected). Regions are ordered sequentially from strongest to weakest negative MvAE value; light gray bars represent the corresponding control MvAE values derived from the resting-state dataset, highlighting the absence of negative adaptation profiles. **(D)** Surface projection of all significant negative MvAE values from panel C mapped onto the cortical mantle using the 300-region Schaefer parcellation, demonstrating robust homeostatic inversion effects distributed prominently across the visual, dorsal attention, and Default Mode Networks (DMN).

In Panel B, we repeated the analysis using matched, segmented resting-state data as a control. Without external stimulation, MvAE values were mostly positive (few between −0.05 < R < 0) and ISC values remained below R = 0.3. This control data derived significance thresholds of R* > 0.25 for ISC and MvAE* < 0 (q(FWER) < 0.01). Crucially, while real movie clips showed a negative correlation, the control analysis revealed a significant positive linear trend between ISC and MvAE (R = 0.3, p<10^-5^).

In Panel C, we addressed the second question of the spread of the MvAE across cortical areas. To that end, all the brain regions that exhibited significant MvAE value from at least one movie-clip, were mapped on the cortical surface. We found 218 regions that showed a significant MvAE effect out of the 254 cortical regions showing significant ISC (R*>0.25, *q*(FWER) <0.01)- i.e. 87% of the regions activated by the movie manifested a significant MvAE. To examine whether the MvAE was associated preferentially with certain brain networks the following analysis was conducted. The average MvAE values was calculated per brain region, only from movie-clips that induced significant negative MvAE, and ordered them from the highest anticorrelation to the lowest. The strongest anticorrelation MvAE values were prevalent in the visual, dorsal attention and DMN areas, as depicted in Figure 5D.

## Discussion

Neural homeostasis, also referred to as rescaling or adaptation, serves as a critical regulatory mechanism that maintains brain activity within an optimal dynamic range, enabling both stability and flexibility in the face of persistent or extreme changes in internal and external conditions. Interestingly, such mechanism seems to be involved in pathological conditions such as neurodegeneration diseases^28^. A substantial body of research has been focused on neural adaptation mechanisms following persistent activations. Well studied effects included motion after-effects^4-6^, light adaptation effects in the retina^29^, or repetition suppression effects, were explored mainly by using artificial generated stimuli, targeting specific brain regions, usually visual areas. Importantly, we are still in the dark concerning whether, to what extent and how widespread such adaptation effects play out during natural, ecological conditions. For example, while engaging in audio-visual movies, which unfold over time. Put simply- it is not known whether neuronal sensitivity remains stable or is dynamically rescaled during ecological, daily life.

Here, using the HCP 7T fMRI dataset for movie-clips, we examined the 20 sec rest periods following these clips to address this question. Lacking sensory stimulations, such rest periods offer a sensitive regime in which traces of adaptation and rescaling produced during the preceding movie clips may be exposed. We observed that the movie clips induced a significant and widely spread adaptation and rescaling effect across many brain regions as depicted in the examples of Figures 3 and 4.

As expected from the adaptation mechanism, regions involved in this adaptation, which we have termed the Movie After-Effect (MvAE) were strongly and persistently affected (activated or in-activated) by the movies. This was directly evident by the high levels of inter-subject synchronization across subjects^21^ of regions eliciting the MvAE. Thus, the MvAE was observed across 218 out of 251 (87%) regions with high ISC (Figure 5c), i.e. regions that were actively involved with movie processing across subjects. This high percentage suggests that the MvAE is a global cortical phenomenon that could be found across the entire cortical mantle as long as cortical regions were highly and persistently involved.

Specifically, and as predicted from a homeostatic mechanism, voxels that showed high activation during the movie reduced their activity below baseline during the post-movie rest, proportionally to their end-of-clip magnitudes, while, conversely, voxels that showed pronounced inactivations tended to increase their activity above baseline during rest. These negative correlations between patterns of voxels activity at the end of clips and patterns of activity towards the end of the resting interval, were robust enough to allow informative MVPA classification of these rest intervals across subjects and brain regions (Figure 2B).

### The Origin of the BOLD MvAE

What drives the BOLD signal below the baseline activity, following high persistent activity? In a seminal work, Shmuel et al.^8^ addressed this question by applying electrical recordings simultaneously with functional MRI (fMRI) in anesthetized macaque monkeys. Their findings demonstrate that the reduction observed in the BOLD fMRI signal following high activation was tightly coupled to a decrease in neuronal activity below spontaneous baseline activity. A possible cellular mechanism that could explain the decrease in firing rate below baseline, after prolong activation was revealed by intracellular recordings of cat primary visual cortical neurons in vivo. These recordings showed that adaptation to a high-contrast stimulus was associated with a hyperpolarization^9^ that was proportional to the reduction of firing rate^10^. This demonstration indicates that neuronal hyperpolarization can provide a plausible mechanism underlying the MvAE found here using BOLD fMRI. It should be noted that such neuronal effects do not necessarily rule out possible contribution of vascular changes to the MvAE as well ^30^, an issue that is still debated^8,31–35^.

### MvAE in Brains and Machines

The MvAE highlights a robust mechanism of on-going adaptation of neural responses during naturalistic stimulation. The dynamic properties of the MvAE strongly point to its functional role in maintaining a stable dynamical range of the neural responses along time, by avoiding activity saturation or undershoot. Intriguingly, a parallel mechanism - batched normalization – has been effectively used in deep neural networks ^36^ in order to facilitate learning, minimize overfitting (by acting as regularization) and covariate shifts. This process is remarkably similar to the adaptation effect reported here in that over-activation of each artificial neuron across mini batches is reduced so as to maintain similar dynamical range of the inputs to all layers. This interesting case of Brain-Machine convergent evolution ^37^ further supports the functional significance of the MvAE reported here even during on-going naturalistic experience.

### Perceptual After Effects

Our findings of adaptation mechanisms observed during post-movie viewing closely parallel those identified in classic laboratory studies of visual adaptation. Importantly, such adaptation has been demonstrated to be associated with perceptual aftereffects. For example, the motion aftereffect—a phenomenon wherein, after sustained exposure to motion in one direction, stationary images are perceived as moving in the opposite direction—is well documented as arising from adaptation in direction-selective neurons within visual cortex ^4^. Similar principles apply to other robust aftereffects, such as the tilt aftereffect, resulting from adaptation of orientation-selective neurons in early visual cortex ^38^, color aftereffect demonstrating adaptation within color-opponent channels ^39^, and size or spatial frequency aftereffects, pointing to the plasticity of feature-selective neural populations ^40^, all of which underscore the broad role of neural adaptation in shaping perception. configuration- related vision exhibits analogous effects, as in the face aftereffect, where exposure to a particular facial expression biases perception of neutral faces ^41,42^.

While much of the literature on aftereffects has focused on low-level stimuli, our results reveal that naturalistic, dynamic stimulation—such as movie viewing—unleashes similar and widespread adaptation effects across the cortex, including attentional networks and some of the DMN regions (Fig 5B). This widespread pattern of neural adaptation extends prior studies, which have largely uncovered such effects in localized and mostly lower-level processing areas (Clifford et al., 2007; Webster, 2011; Anstis et al., 1998). Our findings thus extend classical frameworks, suggesting that adaptation and rescaling serve as a ubiquitous mechanism underlying optimal functional processing across multiple brain systems and during rich, naturalistic perception.

Our results raise the intriguing possibility that even during daily, ecological life- our cortical synaptic networks’ weights may not remain fixed but rather be dynamically renormalized leading to ongoing perceptual and cognitive aftereffects. In analogy with low-level adaptation, such effects should potentially lead to long-lasting perceptual modulations. However, it appears that there is a notable, gap in the literature regarding research into the perceptual consequences of prolonged exposure to complex, dynamic naturalistic environments such as movies. Are viewers also subject to subtle color shifts, distortions in perceived size, or biases in orientation judgments after watching a film? Only a few recent studies have addressed this issue- and indeed reported specific perceptual recalibrations, such as altered speed judgments after adaptation to slow- or fast-motion video ^43^.

However, the widespread nature of the MvAE suggests that the scope of post-movie perceptual changes may extend beyond these low-level visual features. For instance, it is plausible that prolonged engagement with a film narrative could influence subsequent high order cognitive processes. Evidence suggests that internally-generated thought processes, such as auditory-narrative thinking, can persist even after the stimulus has ended ^44^, raising the possibility that our observed post-movie effects might be linked to lingering cognitive activity. Whether the adaptation effects that we observed during post-movie resting intervals might be related to an equivalent “adaptation” of cognitive processes such as short-term memory traces or play other important aspects of information processing related to prior, persistent, activations, remains to be explored.

## Methods

### Experimental Procedures

#### Subjects

From the initial 184 subjects in the HCP S1200 7T release, data from 170 healthy young adults were included in the movie-watching analysis. For the resting-state control analysis, we used the same subjects, resulting in a cohort of 170 subjects.

subjects were partitioned into 17 independent, non-overlapping groups of 10 subjects each, to enhance the signal-to-noise ratio (SNR) and stabilize neural patterns, for both movie-watching and resting-state control analyses. This ‘super-subject’ aggregation strategy creates a robust group-level representation for multivariate analysis.

#### MRI data acquisition

All fMRI data were acquired on a Siemens 7 Tesla Magnetom scanner as part of the Human Connectome Project (HCP) S1200 release. Each participant completed four scan sessions over two to three days. Our analysis focused on the first and last sessions, which included the movie-watching and adjacent resting-state runs.

Functional runs were acquired using a gradient-echo echo-planar imaging (EPI) sequence with the following parameters: TR = 1000 ms, TE = 22.2 ms, flip angle = 45°, field of view = 208 × 208 mm, matrix = 130 × 130, voxel size = 1.6 mm³, number of slices = 85, multiband factor = 5, iPAT = 2, echo spacing = 0.64 ms, partial Fourier = 7/8, bandwidth = 1924 Hz/Px. Phase encoding direction alternated between sessions: PA for REST1, MOVIE2, MOVIE3; AP for REST4, MOVIE1, MOVIE4.

#### Stimuli and experimental design

Each session began with a resting-state run (REST1 or REST4), followed by two movie runs (MOVIE1/2 and MOVIE3/4). During resting-state runs, participants fixated on a crosshair. During movie runs, they passively viewed 4–5 audiovisual clips interleaved with 20-second resting intervals (black screen with the word “REST”).

The full stimulus set included 18 movie clips, including repeated test-retest segments, shown in fixed order across all subjects. Two runs featured Creative Commons independent films (MOVIE1, MOVIE3) and two runs included Hollywood clips (MOVIE2, MOVIE4). The final clip in each run was a test-retest montage repeated identically across runs. Each clip lasted between 63–258 seconds, and all clips were accompanied by synchronized audio via Sensimetric earbuds.

A detailed mapping of clip timing and run organization is shown below: clip index, clip name, and clip onset/offset (in TR units) are as follows: Each clip was followed by an exactly 20-TR (20-second) rest interval. A separate session consisting of continuous rest (“full resting state”) was also acquired for all subjects. A total of 59,412 grayordinate voxels (Grayordinates space) were analyzed, mapped into 300 cortical ROIs based on the Schaefer-300 parcellation^27^, and aligned to 7 functional networks.

The final test-retest clip in each movie run was excluded from analysis because it was repeated identically across runs and thus unsuitable for modeling unique stimulus responses. Below is a table showing the clip schedule (excluding test-retest clips), including clip ID, run name, position in run, short name, and temporal boundaries in TRs. Each Movie scan is initiated with 20 seconds of rest before the movie begins.

| clip_id | run | clip_in_run | name | start_TR | stop_TR |
| --- | --- | --- | --- | --- | --- |
| 1 | MOVIE1_7T_AP | 1 | twomen | 20.0 | 264.04 |
| 2 | MOVIE1_7T_AP | 2 | bridgeville | 284.08 | 505.70 |
| 3 | MOVIE1_7T_AP | 3 | pockets | 525.75 | 713.75 |
| 4 | MOVIE1_7T_AP | 4 | overcome | 733.79 | 797.54 |
| 5 | MOVIE2_7T_PA | 1 | inception | 20.0 | 246.70 |
| 6 | MOVIE2_7T_PA | 2 | socialnet | 266.75 | 525.33 |
| 7 | MOVIE2_7T_PA | 3 | oceans | 545.37 | 794.58 |
| 8 | MOVIE3_7T_PA | 1 | flower | 20.0 | 200.54 |
| 9 | MOVIE3_7T_PA | 2 | hotel | 220.58 | 405.08 |
| 10 | MOVIE3_7T_PA | 3 | garden | 425.12 | 629.20 |
| 11 | MOVIE3_7T_PA | 4 | dreary | 649.25 | 791.75 |
| 12 | MOVIE4_7T_AP | 1 | homealone | 20.0 | 252.29 |
| 13 | MOVIE4_7T_AP | 2 | brokovich | 272.33 | 502.16 |
| 14 | MOVIE4_7T_AP | 3 | starwars | 522.20 | 777.37 |

#### Preprocessing

Functional MRI data were obtained from the Human Connectome Project (HCP) Young Adult dataset and were processed using the HCP minimal preprocessing pipelines ^25^. Briefly, functional images underwent gradient-distortion correction, motion correction, and susceptibility-induced distortion correction using reversed phase-encoding acquisitions. Functional images were registered to each subject’s structural T1- weighted image using boundary-based registration and resampled to standard grayordinate space. Cortical time series were projected onto the cortical surface using ribbon-constrained volume-to-surface mapping, while subcortical signals were retained in volumetric form and combined with surface data into CIFTI dense time-series (.dtseries.nii) files. Cross-subject surface alignment was performed using the MSMAll multimodal surface matching algorithm^45,46^. The time series were high-pass filtered (cutoff = 2000 s) and denoised using ICA-FIX, an automated independent component analysis–based artifact removal method that identifies and regresses out components associated with motion, physiological noise, and scanner artifacts^47,48^. The resulting cleaned grayordinate time series (*_Atlas_MSMAll_hp2000_clean.dtseries.nii) were used for subsequent analyses, for both movie-watching and resting-state data.

### Data Analysis

#### Parcellation — ROI Definition & Feature Dimensionality

Following preprocessing, the data in CIFTI grayordinate space consisted of 59,412 cortical grayordinates. For spatial segmentation, we used the Schaefer-300 parcellation with a 7-network solution^27^, which divides the cortex into 300 functionally defined ROIs based on intrinsic functional connectivity. Each voxel was assigned to one ROI, resulting in 83–428 voxels per parcel (median = 152). Rather than averaging voxel time series within ROIs, we retained the full voxel-level signal for each subject and parcel to preserve spatial heterogeneity within regions. This allowed us to later assess whether observed effects manifested uniformly or focally within regions

We define the voxel-level time series matrix for each subject (s) and region (roi) as:

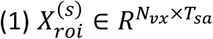

Where 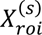 is the BOLD time series matrix for subject (s) and region (*roi)*, *N_vx_* is the number of voxels in the ROI, *T_sa_* is the number of timepoints in the scan.

This voxel-level representation preserves intra-parcel heterogeneity until later analytic stages, enabling us to test whether normalization manifests uniformly or focally within ROIs.

### Mean-Norm-Base Normalization

Voxel Normalization of BOLD signals is critical in naturalistic paradigms, in particular for classification paradigms. To prevent ‘temporal leakage’ during the decoding of post-stimulus resting intervals, voxel normalization was anchored to the initial 20-second unstimulated rest period of each scan. We calculated the Percent Signal Change (*PSC*) relative to this baseline

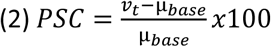

where *v_t_* is the BOLD activation of voxel *v* at time *t*, during a movie-clip, and *μ_base_*is the voxel-wise (*v_t_*) mean during the first 20-sec of the scan related to this movie-clip.

### Subject Grouping

To increase signal-to-noise ratio (SNR), we grouped participants into 17 non-overlapping groups (10 subjects each) and averaged their normalized matrices.

We tested the stability of this grouping by randomly shuffling group assignments 50 times. Classification accuracy varied by <1.5% across permutations, demonstrating robustness to grouping variability. All machine learning analyses employ leave-one-group-out (LOGO) cross-validation, ensuring no information leakage.

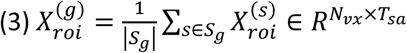

Where 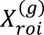 is the group-averaged normalized matrix for group (*g*), *S_g_* is the set of subjects in group, *N_vx_* is the number of voxels per region *roi,* and *T_sa_* is the number of timepoints in the scan.

#### Resting-State Data — Control Analysis

In addition to the movie-watching sessions interleaved with short resting intervals, participants also completed a separate scan consisting entirely of resting-state data. During this session, subjects lay still in the scanner while viewing a black screen with no auditory or visual stimulation. This “full resting state” scan provides an extended baseline that is not confounded by preceding or anticipated stimuli. This condition was critical for evaluating the specificity of our observed effects. By comparing neural activity during movie-adjacent rest with activity from this continuous rest period, we could determine whether observed after-effects were driven by stimulus-related neural adaptation rather than spontaneous fluctuations or scanner drift. In our pipeline, full rest data was subjected to the same subjects, preprocessing and parcellation steps and used to establish a global control condition in multiple analyses, including validation of the normalization procedure and estimation of baseline variability.

### Temporal Feature Construction

#### Movie-clip features

We used the voxel pattern of the last 5-sec of each movie-clip from a local brain region (out of 300 regions), as a signature of that clip. We averaged the BOLD signal of the last 5 TRs of each clip, and across 10 subjects per group, producing a single high-SNR vector per group, per clip, per region.

#### post-clip resting-state features

For the rest periods following each of the movie-clips, we slide a 5-TR window over the 20-TR interval (stride = 1), yielding 14 windows per rest block. We averaged the BOLD signal of the last 5 TRs of each clip, per window, and across 10 subjects per group, producing a single high-SNR vector of voxels per region, per group, per clip. This allowed us to track temporal changes in voxel patterns post-stimulation.

#### Movie Temporal Feature Mean

We average the last **5 TRs** of each clip (per region) to obtain a high-SNR snapshot:

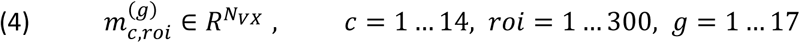

Where 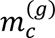 is the group-averaged BOLD vector of length *N_VX_*, the number of voxels of brain region *roi*

,at the end of movie clip *C*. Slide a 5-TR window (stride 1 TR) over the 20-TR rest, yielding **14** windows both for rest between movie and movie matrices.

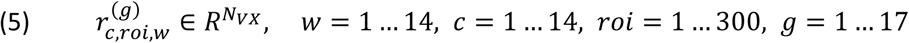

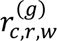 is the BOLD vector for the *w* window of the rest that follows clip (*c*), with dimensionality *N*_vx_, the total number of voxels from region *roi*.

### Movie After-Effect (MvAE)

Briefly, MvAE value is the correlation value that was calculated for each brain region, per clip, per group of subjects (17 groups of 10 subjects each), by correlating between voxels pattern from the end of a movie-clip with voxels-pattern from the last window of rests intervals (Figure 1A, win14) following a movie-clip. Hence, MvAE value is a measure for adaptation driven by movie-clips.

Overall, we computed the Pearson correlation between the last 5-sec of the movie-clips voxel pattern and each of the voxel pattern from the 14 sliding rest windows (w) within the same ROI, clip, and group. This yielded a 14-point trajectory per ROI, reflecting how the adaptation evolved during rest.

Note – the ***MvAE value*** is only the correlation with the last window of the rest period followed by a movie-clip.

For each (clip *c*, group *g*, region *roi, window w*), we compute the Pearson correlation.

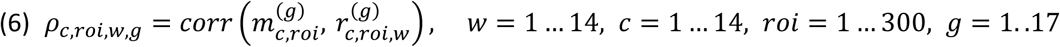

Where *ρ_c_*_,*roi*,*w*,*g*_ is the Pearson correlation between the movie-clip voxel pattern and the rest window (*w*), for clip (c), brain region (*roi)*, and group (*g*). The vectors are restricted to the voxel set inside (roi).

Group-averaging and stacking clips form the trajectory matrix:

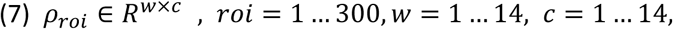

Where *ρ_roi_* is the group-averaged correlation matrix for region (roi), with 14 rows for 14 rest windows

and 14 columns for 14 movie clips (c). The MvAE values presented in the figure 4 are for w=14 (the last 5-sec rest window following a movie-clip).

### SVM Classification — voxel wise Decoding of Clip Identity

We trained linear-kernel SVMs to classify clip identity based on either movie or rest voxels patterns within each ROI. For movies, we stacked 14 clip vectors (voxel pattern for the last 5-sec of the clip) across 17 groups. For each ROI, the SVM utilized a design matrix consisting of **238 samples (14 movie clips across 17 independent groups)**, where the number of features corresponded to the total voxel count within that specific region. When analyzing the resting-state sliding windows, the dataset was expanded to include 14 temporal snapshots per clip, providing a high-resolution temporal profile for the decoding analysis. To preserve the fine-grained spatial topography and functional heterogeneity within each cortical parcel, all analyses were performed on the full voxel-level representation without dimensionality reduction. The classification values that we present is for the last window of the rest interval.

**Movie-clip decoding** uses rest-based design matrix:

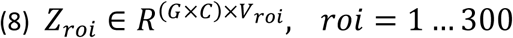

Where *Z_roi_* is the design matrix for *roi*, stacking 14 movie feature (c) vectors across (G=17) groups. Each row has *V_roi_* features (voxels in *roi*, the average of the last 5-sec of the movie-clip).A linear-kernel SVM (C = 1.0) is trained with the **LOGO-CV** (test fold = 14 samples). The final result of this process is a 300- element accuracy vector across the brain, where each element represents the classification accuracy for one ROI.

**Rest decoding** uses rest-based design matrix:

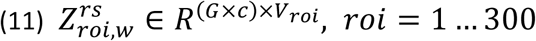

Where 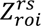 is the rest-based design matrix, for each rest window (w) for 14 clips (c) across 17 groups (G). Each row contains the voxel-wise activity pattern for a single window, with *V_roi_* features (voxels in region roi). Balanced classes ensure chance = 1/14. No dimensionality reduction is applied; later correlation analyses validate that high accuracies coincide with interpretable after-effect signatures, thereby mitigating concerns about overfitting.

#### Inter-Subject Correlation (ISC)

ISC was calculated by correlating group-level activation patterns across all pairs of groups for each movie clip ( the last 5-sec of each movie clip was averaged. ISC was employed on the spatial(voxel) pattern across subjects). The resulting 17×17 matrix was Fisher-z transformed, averaged (lower triangle), and inverse-z transformed to yield a single ISC value per clip. This was repeated at the parcel level.

### Resting-State Data — Control Analysis

In addition to the movie-watching sessions interleaved with short resting intervals, participants also completed a separate scan consisting entirely of resting-state data. During this session, subjects lay still in the scanner while viewing a black screen with no auditory or visual stimulation. This “full resting state” scan provides an extended baseline that is not confounded by preceding or anticipated stimuli. This condition was critical for evaluating the specificity of our observed effects. By comparing neural activity during movie-adjacent rest with activity from this continuous rest period, we could determine whether observed after-effects were driven by stimulus-related neural adaptation rather than spontaneous fluctuations or scanner drift. In our pipeline, full rest data was subjected to the same subjects, preprocessing and parcellation steps and used to establish a global control condition in multiple analyses, including validation of the normalization procedure and estimation of baseline variability.

### Statistical and control analysis

To test whether stimulus-induced processing of the movie-clip predicted post-stimulus rest adaptation, and no other factors, we repeated the exact same analysis, in order to get the null correlations for the ISC and MvAE, but using the full resting-state data from the same subjects, segmented randomly into the lengths of movie-clips and resting-states intervals in between movie-clips, 1000 times. Fo each iteration we kept the maximum ISC across the entire brain and the minimum MvAE across the entire brain. We controlled the FWER by defining a threshold (R*) at the q*100th percentile of the null distribution across the whole brain, of maximum (ISC) and minimum values (MvAE). only correlation with correlation value (*R*) above (and below) the threshold derived from null distribution (*R*\*) were considered significant after correction for multiple comparisons. The thresholds for each condition were as follows (all for *q*<0.01): ISC *R\**=0.25; MvAE *R\**=0.

## Data and code availability

The movie-watching and resting-state fMRI data used in this manuscript are part of the publicly available and anonymized HCP database (https://www.humanconnectome.org).

Any additional information required to reanalyze the data reported in this paper is available from the corresponding author upon request.

## Funding Statements

The authors declare no funding was received for this work.

## Contributions

Conceptualization and writing by E.S., R.M., and N.Y. Supervision by E.S. and R.M.; Methodology, project administration, and code programming by N.Y. and E.S.; Data curation, formal analysis, investigation, validation, and visualization by N.Y and E.S.

## Competing interests

The authors declare no competing interests.

## Notes

### Competing Interest Statement

The authors have declared no competing interest.

